# SPRTN Protease and Checkpoint Kinase 1 Cross-Activation Loop Safeguards DNA Replication

**DOI:** 10.1101/458026

**Authors:** Swagata Halder, Ignacio Torrecilla, Martin D. Burkhalter, Marta Popovic, John Fielden, Bruno Vaz, Judith Oehler, Domenic Pilger, Davor Lessel, Katherine Wiseman, Abhay Narayan Singh, Iolanda Vendrell, Roman Fischer, Melanie Philipp, Kristijan Ramadan

## Abstract

The SPRTN metalloprotease is essential for DNA-protein crosslink (DPC) repair and DNA replication in vertebrate cells. Cells deficient in SPRTN protease activity exhibit severe DPC-induced replication stress and genome instability, manifesting as premature ageing and liver cancer in humans and mice. Strikingly, SPRTN-deficient cells also show a severe G2/M checkpoint defect and fail to activate a robust checkpoint kinase 1 (CHK1) signalling cascade normally triggered in response to replication stress. Here, we show that SPRTN activates the CHK1 signalling cascade during physiological (steady-state) DNA replication by proteolysis-dependent eviction of CHK1 from chromatin. The N-terminal CHK1 fragments cleaved by SPRTN protease still possess kinase activity and phosphorylate SPRTN. This CHK1-dependent phosphorylation stimulates SPRTN recruitment to chromatin to further promote CHK1 eviction from chromatin, DPC removal and unperturbed DNA replication. Our data suggest that a SPRTN-CHK1 cross-activation loop is essential for steady-state DNA replication and protection from DNA replication stress, a major source of genome instability in diseases that cause premature ageing and cancer. In addition, we disclose a mechanism of CHK1 activation during physiological DNA replication, when long stretches of single stranded (ss) DNA that are induced by genotoxic stress and activate canonical and robust CHK1 signalling are not present.

## Introduction

The timely completion of DNA replication is essential for genome integrity and preventing the onset of cancer, premature ageing and developmental disorders ^1,2^. DNA replication is constantly threatened by many factors, including DNA lesions, collisions with the transcription machinery and repetitive DNA sequences. Cells have evolved robust DNA damage tolerance and DNA damage response pathways to cope with the different lesions and obstacles that challenge the progression of DNA replication forks ^3–6^. In particular, the ATR (Ataxia telangiectasia Rad3 related)-CHK1 signalling cascade is the major regulator of the response to replication stress ^7,8^. This cascade performs multiple functions in response to DNA replication stress, including regulating DNA replication origin firing, stabilising stalled replication forks, and delaying mitotic entry by preventing CDK1/2 hyper-activation ^5,9^

The main stimulus for CHK1 activation is replication protein A (RPA)-coated single-stranded (ss) DNA that typically forms upon DNA replication fork stalling due to uncoupling of the Cdc45-Mcm2-7-GINS (CMG) helicase complex from DNA polymerases ^5^. This ssDNA-protein structure recruits the ATR-ATRIP protein kinase complex which activates CHK1 by phosphorylating serines 317 and 345 to expose the catalytic N-terminal domain of CHK1 ^10^. Upon phosphorylation, CHK1 is released from chromatin by an as yet unknown mechanism and spread throughout the nucleus and cytoplasm to regulate the activity of its substrates and safeguard genome stability ^9,11,12^. The CHK1 signalling pathway can also be activated in S-phase by ssDNA generated after 5’-3’ end resection of double strand DNA breaks (DSB) ^13^.

However, given that CHK1 activity is required for physiological DNA replication fork progression - when long stretches of ssDNA are scarce - the question of how CHK1 is activated during steady-state DNA synthesis remains unanswered ^14–18^. Identifying the mechanisms that regulate CHK1 signalling under physiological conditions is therefore essential to understand how cells survive and preserve genomic stability ^19^.

We and others have recently identified the SPRTN metalloprotease as being a constitutive component of the DNA replication machinery and essential for DNA replication ^20–24^. The essential role of SPRTN in safeguarding genome stability is demonstrated both in human disease and in animal models. Monogenic, biallelic *SPRTN* germline mutations cause Ruijs-Aalfs Syndrome (RJALS), a rare disease characterised by genomic instability, premature ageing and hepatocellular carcinoma ^20,25,26^. Furthermore, SPRTN haploinsufficient mice develop RJALS-like phenotypes, while complete SPRTN knockout is embryonic lethal ^27^. SPRTN has recently been identified as an essential core fitness gene in humans ^27–30^. Finally, downregulation of SPRTN in zebrafish severely impairs normal embryonic development and increases embryonic lethality ^25^.

The source of genome instability in both RJALS patient cells and SPRTN-deficient human and mouse cells was recently demonstrated to arise from replication stress caused by the accumulation of replication-blocking DNA-protein crosslinks (DPC) ^20,22,23^. DPCs are formed by various aldehydes including the well-known DPC inducing agent formaldehyde (FA), which are by-products of metabolic processes such as lipid peroxidation or histone and DNA demethylation ^31–33^. As SPRTN protease activity is required to cleave DPCs, defective SPRTN protease activity results in profound replication stress, visualised as increased fork stalling and significantly reduced DNA replication fork velocity. Strikingly, we observed a severe G2/M-checkpoint defect in SPRTN-deficient cells treated with genotoxic agents that interfere with DNA replication ^25^. The G2/M checkpoint was, however, completely functional after the induction of non-replication-associated DNA strand breaks using ionising radiation, suggesting that SPRTN-defective cells lack the ability to activate CHK1 in response to replication stress when replication forks are still intact ^25^.

Here, we demonstrate that SPRTN stimulates CHK1 function during physiological DNA replication and vice versa. SPRTN proteolytic activity (proteolysis) evicts CHK1 from replicative chromatin/sites of DNA replication what allows physiological CHK1 function during steady-state DNA replication. We also show that SPRTN proteolysis cleaves the regulatory/inhibitory C-terminal domain of CHK1 *in vitro* and *in vivo*, and thus releases active N-terminal CHK1 products. These N-terminal CHK1 products, when ectopically expressed, are sufficient to stabilise DNA replication forks, rescue embryonic development and genome stability that are compromised by depletion of SPRTN to approximately 30% of wildtype cells. This rescue is dependent of CHK1 phosphoryalting the C-terminus of the residual SPRTN, further promoting SPRTN recruitment to chromatin for the removal of DPCs in front of DNA replication forks. In summary, we show that a SPRTN-CHK1 crossactivation loop is essential for steady-state DNA replication, embryonic development and survival and is evolutionarily conserved in vertebrates.

## Results

### SPRTN deficiency leads to aberrant CHK1 activity

Analysis of RJALS patient and SPRTN-depleted cells revealed that SPRTN protease activity is essential for DNA replication fork progression, cell cycle progression, and G2/M checkpoint activation after DNA replication stress but not ionising radiation ^20,23,25^. These results suggest that SPRTN bridges DNA replication and G2/M-checkpoint regulation. To re-evaluate these findings, we performed analysis of DNA replication using the DNA fiber assay (Fig. 1a). Depletion of SPRTN by three independent siRNA sequences caused severe DNA replication stress in human embryonic kidney 293 (HEK293) cells, visualised as a reduction in DNA replication fork velocity and an increased frequency of fork stalling (Fig. 1b-d). Short treatment of control cells with a low dose of hydroxyurea (HU), a drug that limits the cellular dNTP pool, was used as a positive control of DNA replication stress phenotypes. In general, as a response to DNA replication stress, cells suppress dormant origin firing, as was visible after HU treatment (Fig. 1e). Interestingly, dormant origin firing was more than 3-fold higher in SPRTN-depleted cells when compared to HU-treated cells. As firing of dormant origins is tightly regulated by the CHK1 kinase ^34,35^ which also controls the G2/M-checkpoint ^36,37^, we asked whether CHK1 signalling was defective in SPRTN-inactivated cells. ATR-CHK1 signalling, visualised by CHK1 S345 phosphorylation, was not activated in SPRTN-depleted cells (Fig. 1f, g) despite the severe DNA replication stress phenotypes (Fig. 1b-e). Accordingly, CHK1 kinase activity was not activated as demonstrated by the lack of CHK1 serine 296 (S296) phosphorylation, the residue that CHK1 auto-phosphorylates once its kinase activity has been stimulated by ATR ^38,39^. Furthermore, despite being in similar cell cycle stages (Supplementary Fig. 1a-c), SPRTN-depleted cells exhibited less CHK1-S296 and -S345 phosphorylation than even unchallenged control cells and ~9-10-fold less compared to HU-treated control cells, which showed similarly reduced replication fork velocity and elevated levels of fork stalling as SPRTN-depleted cells (Fig. 1f, g). Similar findings were observed in two other human cells lines: HeLa (Fig. 3b, c) and U2OS (data not shown). These results show that SPRTN-inactivated human cells fail to activate a robust CHK1 response despite exhibiting severe replication stress phenotypes that would ordinarily be expected to elicit such a response, namely CHK1 chromatin eviction and consequent activation of CHK1 signalling ^11^.

**Figure 1.**
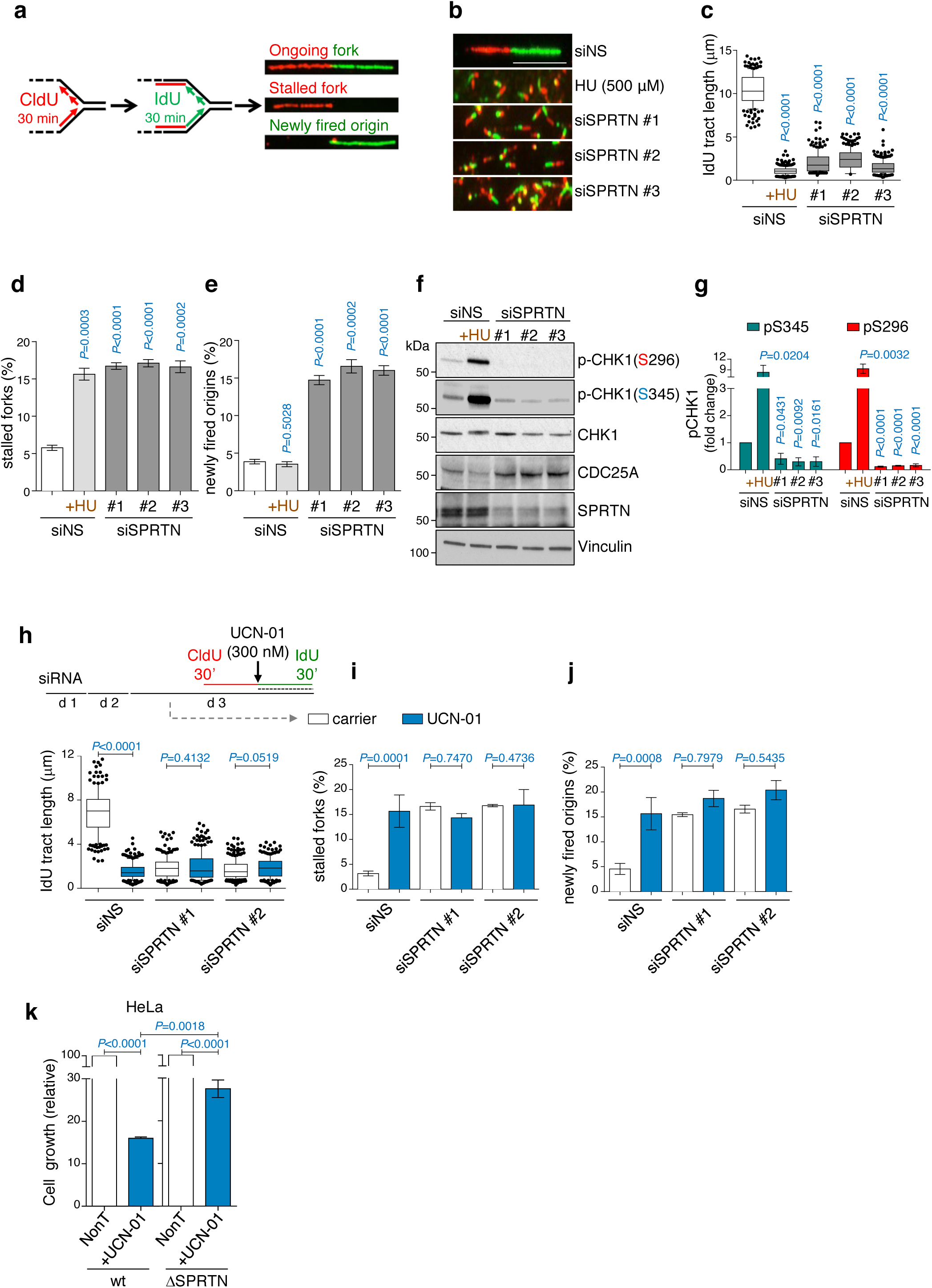
SPRTN depleted cells endure severe DNA replication stress but fail to activate a CHK1 response. **a**, Schematic representation of DNA fiber assay. See also Methods. **b**, DNA fibers obtained from HEK293 cells that have been treated with the indicated siRNAs against SPRTN (siSPRTN) or with hydroxyurea (HU). Scale bar: 10 μm. Data shown are representative images of three independent experiments. **c-e**, DNA fiber assay analysis of **b** showing replication fork length (**c**), stalled replication forks (**d**) and newly fired origins (**e**). **c**, siSPRTN cells exhibit decreased fork velocity. >100 individual IdU tracts were measured per experiment per condition. Data are shown as mean with 25-75% percentile range (box) and 10-90% percentile (whiskers); n=3 independent experiments, two-tailed Student’s *t-* test. **d**, siSPRTN cells undergo increased stalled replication forks. >400 forks were scored per condition per experiment. Mean ± SEM; n=3 independent experiments, two-tailed Student’s /-test. **e**, siSPRTN cells undergo increased newly firing of dormant origin, contrary to cells treated with HU. >400 forks were scored per condition per experiment. Mean ± SEM; n=3 independent experiments, two-tailed Student’s t-test. **f**, Knock-down of SPRTN in HEK293 cells diminishes the phosphorylation status of CHK1 residues S296 and S345, and enhances CDC25A stability, contrary to treatment with HU. Immunoblots were run with whole cell lysates and represent three independent experiments. **g**, Quantifications for **f** of CHK1 phosphorylation signal at S345 and S296 residues, normalised to vinculin. Mean ± SEM; n=3 independent experiments, two-tailed Student’s t-test. **h-j**, Inhibition of cells with the CHK1 inhibitor UCN-01 induces severe replication stress to a similar extent as SPRTN-inactivation, detected by DNA fiber assay analysis. Schematic representation of experimental layout showing addition of UCN-01 with IdU is also shown in **h** (top); d: day. **k**, Cell growth of *wt* and SPRTN-deficient (DSPRTN) HeLa cells in response to the CHK1 inhibitor UCN-01 (300nM) on day 4 after seeding cells at same density (day 0). Mean ± SEM; n=3 independent experiments, two-tailed Students′s *t*-test.

Indeed, SPRTN-inactivated cells accumulate over two-fold more CHK1 on chromatin, similar to cells treated with the DNA-protein crosslinking agent formaldehyde (FA), but opposite to cells treated with HU, where CHK1 is released from chromatin (Supplementary Fig. 1d, e).

To further validate this observation, we treated HEK293 cells with UCN-01, a well-characterised CHK1 inhibitor ^15,40,41^. UCN-01 treatment in control cells caused a severe reduction in replication fork velocity and increased the frequency of new (dormant) origin firing and fork stalling (Fig. 1h-j). UCN-01, however, had no additive effect on replication fork velocity, new origin firing, or fork stalling in SPRTN-depleted cells. In addition, in comparison to wt cells, SPRTN-haploinsufficient HeLa (DSPRTN) cells were relatively but significantly less sensitive to UCN-01 treatment (Fig. 1k). This further suggests that CHK1 is not fully activated in SPRTN-defective human cells. Altogether, these results highlight the importance of CHK1 activity during steady-state DNA replication ^14,15,18^, demonstrate an epistatic relationship between SPRTN and CHK1, and reveal a severe defect in CHK1 kinase activation in SPRTN-defective cells despite them suffering from severe DNA replication stress. We concluded that SPRTN-defective cells lack the optimal (physiological) CHK1 activity required for DNA replication and genome stability, most probably due to their inefficiency in evicting CHK1 from chromatin, and thus activating a steady-state CHK1 signalling cascade.

### SPRTN regulates CHK1 signalling pathway under physiological conditions

To test this conclusion, we investigated the signalling pathway downstream of CHK1 by monitoring the total levels of the CHK1 target, protein phosphatase Cdc25A ^3^. Upon phosphorylation by CHK1, Cdc25A is degraded by the proteasome, as observed in cells treated with HU (Fig. 1f and Supplementary Fig. 1f). Consequently, CDK1/2 are hyper-phosphorylated and become inactive, which leads to intra S-phase and G2/M-checkpoint activation and cell cycle arrest ^42,43^. CDK1 and 2 drive S-phase progression and the G2/M cell cycle transition, but hyper-activation of CDK1/2 negatively influences DNA replication fork stability and causes premature mitotic entry ^44,45^. Hence, the ATR-CHK1-Cdc25-CDK1/2 pathway is necessary to regulate cell cycle progression during DNA synthesis. Due to faulty CHK1 activation, SPRTN-depleted cells hyper-accumulated Cdc25A (Fig.1f and Supplementary Fig. 1f), which in turn dephosphorylates and hyper-activates CDK1/2, as was visible by the increased phosphorylation of total CDK1/2 substrates in HEK293 cell extracts (Supplementary Fig. 1g, h).

### CHK1 overexpression corrects DNA replication stress and genome instability in SPRTN-defective cell

To assess whether the failure to activate CHK1 signalling could explain the DNA replication phenotypes and G2/M defects observed in SPRTN-depleted cells (Fig. 1) ^25^, we ectopically expressed CHK1-wild type (wt) or its phosphorylation (phospho)-defective variants (CHK1-S317A or CHK1-S345A) (Fig. 2a, b and Supplementary Fig. 2a). CHK1-wt, but not CHK1-S317A or CHK1-S345A, restored DNA replication fork velocity, suppressed new origin firing and rescued replication fork stalling in SPRTN-depleted cells. Moreover, ectopic expression of CHK1-wt in SPRTN-deficient cells also corrected chromosomal instability, measured by the number of chromosomal aberrations on mitotic chromosomes (Fig. 2c and Supplementary Fig. 2b). These results further support our initial observation that SPRTN-inactivation leads to an impaired CHK1 signalling cascade, resulting in severe DNA replication stress, a defective G2/M checkpoint and the accumulation of chromosomal aberrations in SPRTN-deficient cells ^25,27^.

**Figure 2.**
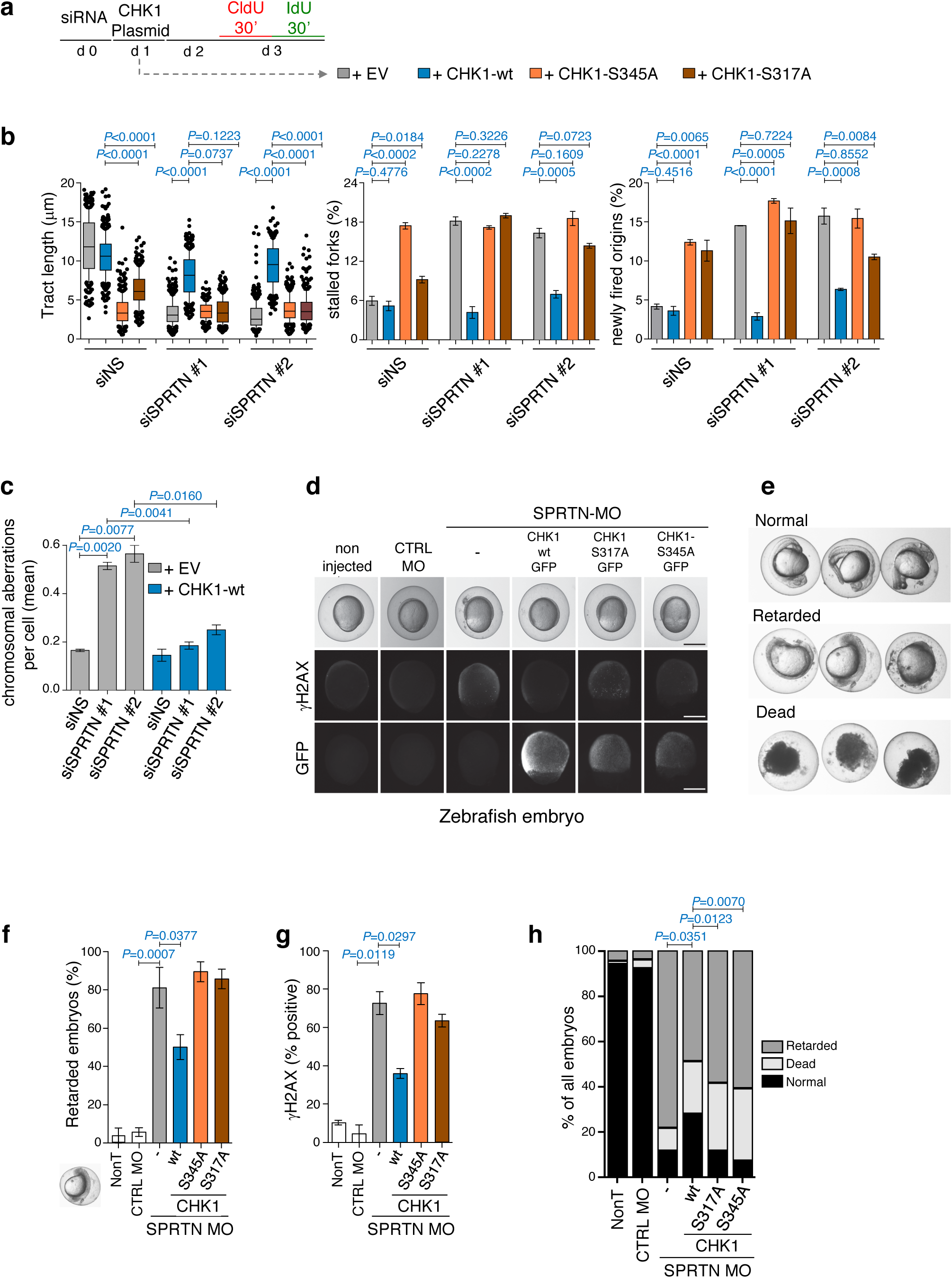
Ectopic CHK1 overexpression rescues SPRTN-deficiency phenotypes in human cell lines and Zebrafish embryos. **a**, Experimental strategy followed to assess the effect of different CHK1 variants on DNA replication by DNA fiber assay (d: day, EV: empty vector). **b**, Rescue of DNA replication defects in SPRTN-depleted HEK293 cells with ectopic expression of CHK1-wt, but not the phospho-deficient CHK1 variants S345A and S317A. Graphs show quantification data from DNA fibers for replication fork velocity (left panel; mean ± 25-75 percentile range (box) ± 10-90 percentile range (whiskers)), newly fired origins (centre panel; mean ± SEM) and stalled replication forks (right panel; mean ± SEM); > 100 DNA fibers were analysed per condition and experiment; n= 3 independent replicates, two-tailed Students′*t*-test. **c**, Overexpression of CHK1-wt diminishes the number of chromosomal aberrations caused by SPRTN knock-down visualized by metaphase spreads from HEK293 cells. >30 cells were scored per condition per experiments; mean ± SEM; n= 3 individual experiments, two-tailed Student′s *t*-test. **d**, Representative images of zebrafish embryos non-injected or injected with either a control or a previously characterised morpholino (MO) against SPRTN at 9-10 hours post fertilization (hpf). To assess CHK1 function, capped RNA encoding different mutants of GFP-tagged human CHK1 were co-injected with the SPRTN MO. Upper row: live images. Middle row: pictures of γH2AX to show DNA damage. Lower row: Pictures of GFP to show expression of CHK1 protein variants. **e**, Representative images for normal, retarded and dead zebrafish embryos at 24 hpf. **f-h**, Overexpression of CHK1-wt, but not of the phospho-deficient CHK1 variants S345A and S317A, rescues developmental defects and DNA damage induced by SPRTN depletion in zebrafish embryos. Data scoring normal, retarded and dead zebrafish embryos are represented in h, whereas percentage of zebrafish embryos that show developmental retardation relative to living embryo are represented in **f**. Mean ± SEM; > 70 embryo were scored in n=4 independent experiments, two-tailed Student′s *t*-test. γH2AX, marker of DNA damage, is represented in g. Mean ± SEM; at least 30 GFP-positive embryos were counted per condition; n=2 replicates, two-tailed Student′s /-test.

### The SPRTN-CHK1 axis is essential for development and genome stability in zebrafish embryos

To investigate and validate our observations so far on an organismal level, we took advantage of the zebrafish model system. We have previously shown that morpholino (MO)-mediated depletion of SPRTN in zebrafish embryos causes severe development defects and accumulation of DNA damage ^25^. When fertilized eggs with SPRTN MO were co-injected with capped RNAs encoding for GFP-CHK1-wt, both developmental retardation and DNA damage (the latter analysed by γH2AX accumulation) were rescued (Fig. 2d-h and Supplementary Fig. 2c). Conversely, reconstitution with RNAs encoding phospho-defective variants of GFP-CHK1, S317A or S345A, failed to rescue the phenotypes of SPRTN depletion in early zebrafish embryos. Altogether, these data suggest that the restoration of CHK1 activity is able to compensate for SPRTN-deficiency in human cells and zebrafish embryos, and that the SPRTN-CHK1 axis is conserved in vertebrates.

### SPRTN does not contribute to CHK1 activation after DNA double strand break formation

Interestingly, when SPRTN-depleted cells were challenged with HU, CHK1 was activated to the same extent as in HU-treated control cells (Fig. 3a, lanes 2, 4, 6 and 8). This CHK1 activation was concomitant with phosphorylation of ATM, CHK2 and RPA, indicating DSB formation ^36,46^. Moreover, the total level of RPA remained unchanged in these cells and phosphorylation of RPA ruled out the possibility that severe replication stress in SPRTN-inactivated cells results in RPA exhaustion ^47^. Similar findings were observed in haploinsufficient HeLa SPRTN cells (Δ-SPRTN) (Fig. 3b, c). Taken together, our data suggest that SPRTN-deficient human cells have a fully functional ATR-CHK1 signalling pathway in the context of DSB repair, but that SPRTN is needed to activate this pathway during physiological/steady-state DNA replication.

**Figure 3.**
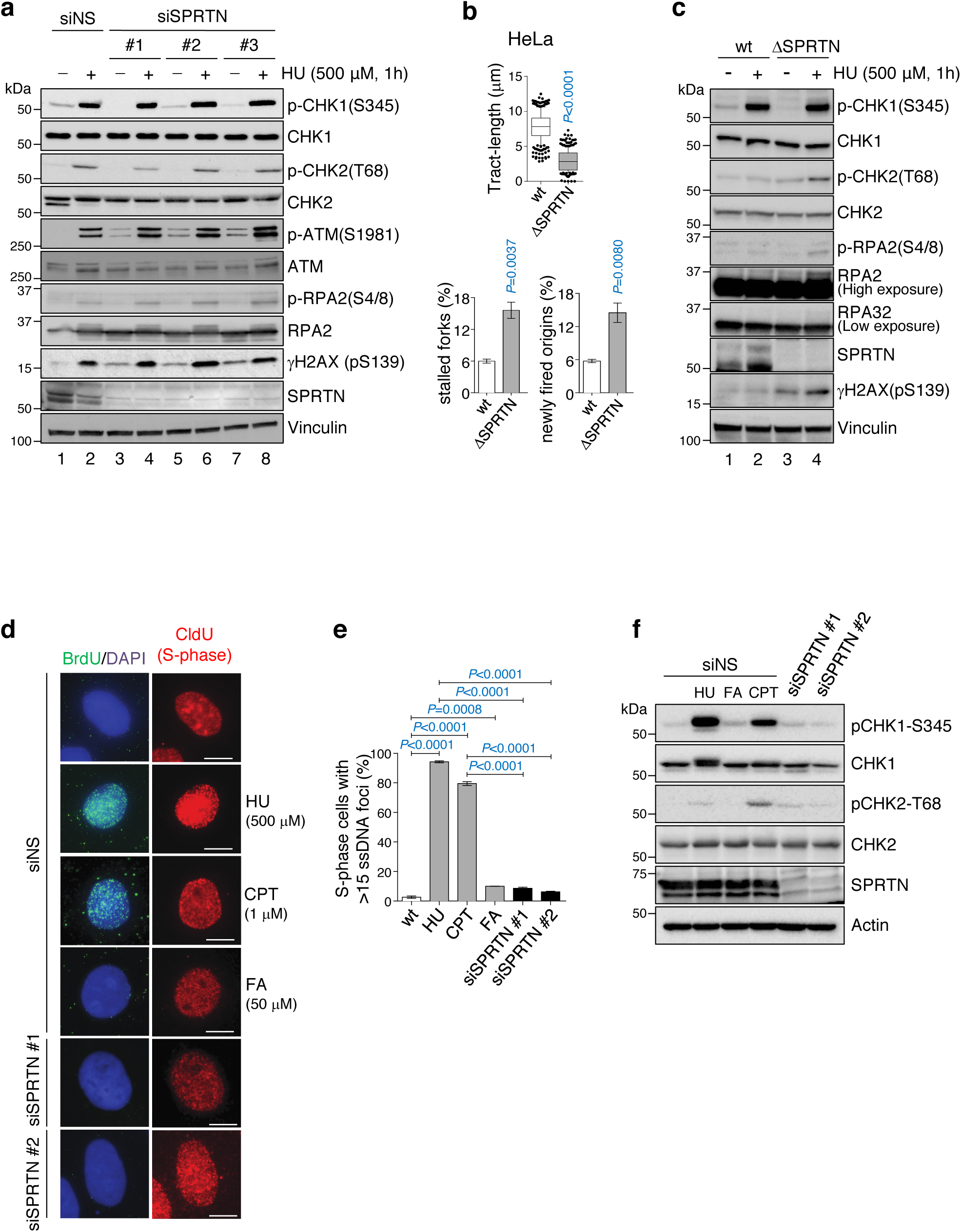
SPRTN does not contribute to activation of CHK1 after DSB formation. **a**, Phosphorylation status of the ATR-CHK1 and ATM-CHK2 pathways in control (siNS) and siRNA SPRTN-depleted HEK293 cells under unchallenged condition or when cells were treated with HU. Whole cell extracts were used for the immunoblots. **b**, DNA fiber assay analysis comparing HeLa-wt *(wild type*, parental) and HeLa-ASPRTN cells: quantification of replication fork velocity (top; mean ± 25-75 percentile range (box) ± 10-90 percentile range (whiskers)), newly fired origins (left; mean ± SEM) and stalled replication forks (right; mean ± SEM). > 100 DNA fibers were analysed per condition and experiment; n=3 experimental replicates, two-tailed Student′s *t*-test. **c**, Analysis of the phosphorylation status of the ATR-CHK1 and ATM-CHK2 pathways comparing Hela-wt and ΔSPRTN under unchallenged or HU-treated conditions. Whole cell extracts were used for the immunoblots. **d-e**, SPRTN-deficient (siSPRTN) or wt cells (siNS) treated with formaldehyde (FA; 50 μM, 1 h) do not exhibit the generation of robust single-stranded DNA foci in S-phase cells (CldU positive). HU (500 μM, 1 h) was used as a positive control for replication stress-induced ssDNA formation; CPT (1 μM, 1 h and 1 h recovery) was used as a positive control for ssDNA induced by DSBs. Data are shown as representative immunofluorescent microscopy images of BrdU foci (ssDNA) in S-phase (CldU positive) HeLa cells (d), and (e) as quantification of cells with more than 15 BrdU foci. >70 individual CldU positive cells were scored per condition per experiment. Mean ± SEM; n=3 experimental replicates, two-tailed Student′s *t*-test. **f**, FA treatment and SPRTN-depletion fail to induce phosphorylation of CHK1 at Ser345 due to the absence of ssDNA formation. HU or CPT were used at the same conditions as in d as positive controls for ssDNA-induced CHK1 and CHK2 activation. Immunoblots of HeLa cells whole cell extracts shown here represent three experimental replicates.

In addition, immunofluorescence-coupled microscopy analysis of ssDNA formation in S-phase cells (CldU positive) by BrdU staining under native conditions further suggested that SPRTN-defective cells do not form extensive ssDNA formation (Fig. 3d-f), a platform for canonical and robust CHK1 activation, despite enduring severe DNA replication stress. Similar results were obtained with FA-treatment. HU treatment was used as a positive control for DNA replication stress-induced ssDNA formation. However, when SPRTN-depleted or FA-treated cells were exposed to a high dose of Camptothecin (CPT) or HU, a striking increase in ssDNA (visualised as BrdU foci), generated by 5’-3’ end resection of DSBs (Supplementary Fig. 3a, b), was observed. This effect ruled out the possibility that this BrdU/ssDNA assay was compromised in SPRTN-deficient cells owing to reduced BrdU incorporation due to defective DNA replication (Supplementary Fig 1a-c). Altogether, these results suggest that SPRTN is essential for activation of physiological CHK1 response during steady-state DNA synthesis, most probably due to removal of bulky DPCs in front of DNA replication forks that prevent CMG helicase progression and consequently ssDNA formation, but dispensable for CHK1 activation after DSB formation, when a huge amount of ssDNA is formed by 5’-3’ end resection.

### SPRTN proteolysis releases CHK1 from replicating chromatin

Given that SPRTN enables DNA replication by proteolysis of its chromatin-bound substrates ^20,22^ and that, in SPRTN-defective cells, CHK1 is not activated during physiological conditions (Fig. 1) but retained on chromatin (Supplementary Figure 1d, e), we wondered whether CHK1 is a substrate of the SPRTN protease. To address this question, we employed four experiments, both *in vitro* and *in vivo*. One, by using cell fractionation we demonstrated that a small amount of CHK1 is constantly bound to chromatin (Fig. 4a, b). Additionally, CHK1 significantly hyper-accumulated on chromatin isolated under stringent and denaturing conditions in SPRTN-inactivated cells during S-phase progression (Fig. 4c, d) suggesting that a small protion of CHK1 is also covalently attached to chromatin. Two, in pull-down experiments using purified SPRTN and CHK1 proteins we demonstrated that these two proteins interact directly *in vitro* (Fig. 4e). Three, by co-immunoprecipitation experiments from HEK293 cells expressing SPRTN-wt or a RJALS patient protease defective variant (SPRTN Y117C), we showed that both SPRTN wt and Y117C form a physical complex with CHK1 and other components of DNA replication machinery such as PCNA and MCM3 ^48^, further suggesting that SPRTN and CHK1 are components of the DNA replication machinery *in vivo* (Fig. 4f). Four, we took advantage of iPOND to demonstrate SPRTN-dependent CHK1 dynamics at replicative chromatin (Fig. 4g, h). iPOND directly demonstrated that both CHK1 and SPRTN are present at/around sites of DNA replication forks (nascent DNA) and travel with the fork as confirmed by their absence from mature chromatin (after the thymidine chase) (Fig. 4h). Importantly, inactivation of SPRTN led to a strong accumulation of CHK1 on both nascent and mature chromatin (Fig. 4i, compare lanes 2 and 3 with 4 and 5).

**Figure 4.**
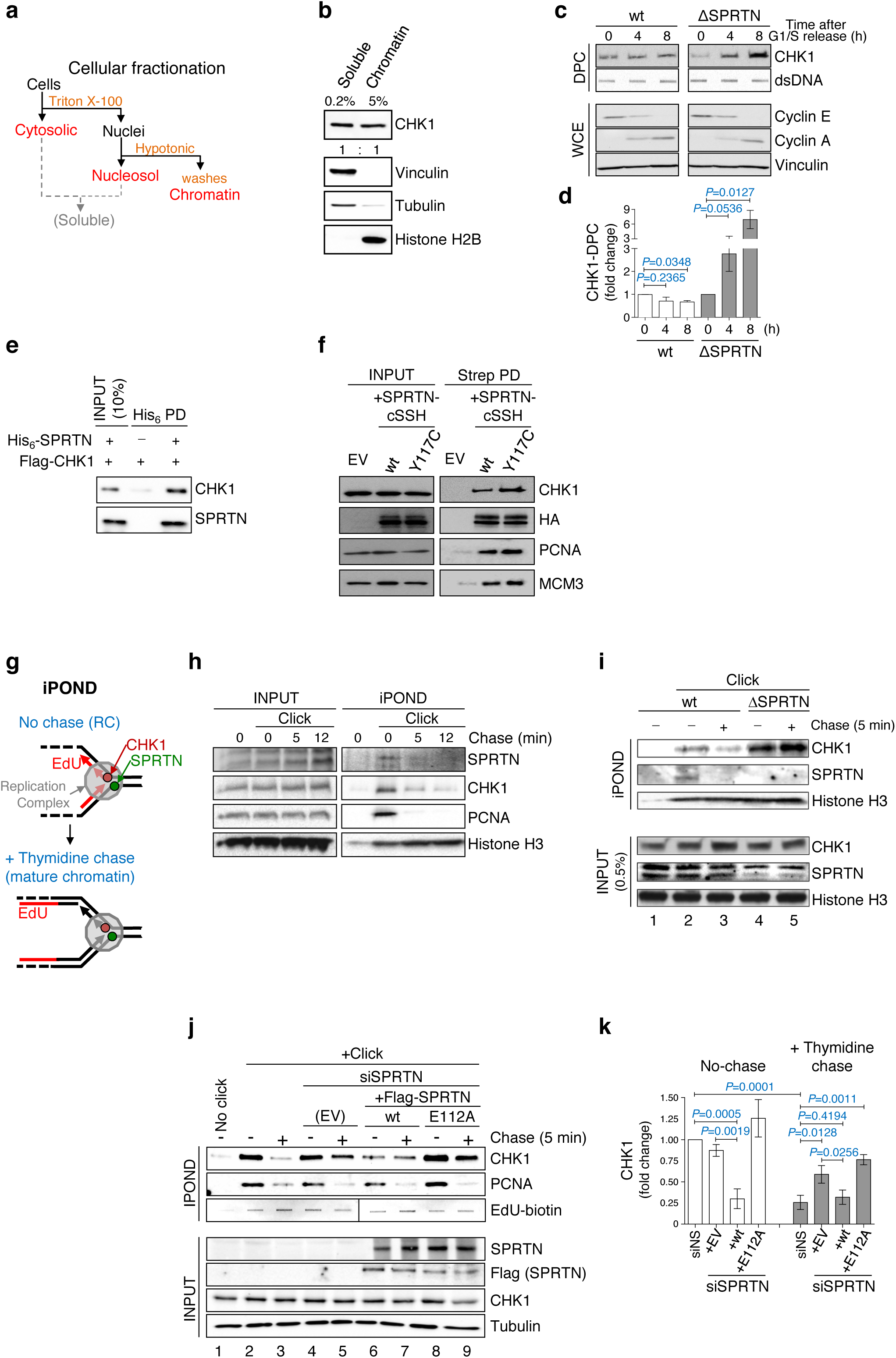
SPRTN protease evicts CHK1 from replicative chromatin. **a**, Diagram of cellular fractionation into cytosolic (soluble), nuclear (soluble) and chromatin (insoluble) fractions. The insoluble fraction was thoroughly cleaned by several washes with 1% NP- 40 and 250 mM NaCl to isolate clean chromatin (see Methods). **b**, Clean chromatin fractionation was confirmed by fractionation protein markers, and it still contains at least 4% of the total cellular pool of CHK1. **c-d**, SPRTN deficiency leads to accumulation of tightly bound CHK1 on chromatin during the S-phase progression. HeLa-wt and HeLa DSPRTN cells were arrested at G1/S boundary by a double thymidine block and then released to progress through S-phase. Upper set: Proteins tightly bound to DNA were isolated by a modified RADAR assay (see Methods) and the total content of CHK1 in these isolates was tested by immunoblotting. Double stranded DNA (ds-DNA) was used as a loading control. Lower set: cyclin E and cyclin A from whole cell extracts were used as cell cycle markers. Data in **d** are the quantification of CHK1 tightly bound to DNA. Mean ± SD; n=3, two-tailed Student′s *t*-test. **e**, SPRTN and CHK1 interact physically *in vitro.* Purified CHK1 was co-precipitated with Ni-NTA from mixtures with recombinant His_6_-tagged SPRTN. Immunoblots represent three replicates. **f**, SPRTN and CHK1 interact physically *in vivo.* CHK1 was co-precipitated with Strep-Tactin sepharose in whole cell extracts lysates of stable Flp-In TRex HEK293 expressing either SPRTN-wt-cSSH (protein fused to Strep-Strep-HA), SPRTN-Y117C-cSSH (Y117C is a SPRTN patient variant with ~ 20% protease activity). Immunoblots represent three replicates. EV: empty vector. **g-h**, iPOND (isolation of proteins on nascent DNA; see Methods) analysis revealed the presence of SPRTN and CHK1 proteins at nascent DNA (click, 0 min) but not on mature chromatin (click after a 4 or 12 min thymidine chase). Cells were treated with EdU for 10 min to label nascent DNA, and then chased (to detect mature DNA) or not with thymidine. PCNA and histone H3 were used as replication fork and loading controls, respectively. Immunoblots represent three replicates. **i**, SPRTN-deficient (ΔSPRTN) HeLa cells exhibit increased retention of CHK1 on mature DNA than SPRTN-proficient (wt) cells. **j-k**, SPRTN protease is necessary for the efficient eviction of CHK1 from replicative chromatin. Expression of SPRTN-wt, but not SPRTN protease inactive variant (E112A), in SPRTN knock-down cells reduced the amount of CHK1 in replicative chromatin and reversed the accumulation of CHK1 in chased chromatin. Mean ± SEM; n=3 experimental replicates, two-tailed Student′s /-test.

In the light of the above observations we used iPOND to directly assess if SPRTN proteolysis plays an essential role in CHK1 release/eviction from sites of (around DNA replication) replicative chromatin. We analysed the abundance of CHK1 by iPOND in SPRTN-depleted cells expressing an empty vector (EV), SPRTN-wt or the SPRTN protease-deficient variant E112A (Fig. 4j, k). Although both SPRTN variants were equally expressed, SPRTN-wt strongly evicted CHK1 from nascent (DNA replication sites) and mature chromatin but SPRTN-E112A did not. A similar effect was observed on total chromatin isolated by biochemical fractionation (Supplementary Figure 4a, b). Altogether, these results directly demonstrate that the SPRTN-CHK1 complex is an integral part of the DNA replication machinery and that SPRTN proteolysis evicts CHK1 from replicating chromatin during DNA synthesis.

### SPRTN protease cleaves CHK1

As SPRTN directly interacts with (Fig. 4e) and evicts CHK1 from sites of DNA replication in a protease-dependent manner (Fig. 4g-k), we investigated whether SPRTN proteolytically cleaves CHK1. Purified Flag-CHK1 was incubated with SPRTN-wt or a protease-deficient variant (E112A) *in vitro* in the presence of dsDNA, an activator of SPRTN metalloprotease activity (Fig. 5a). SPRTN-wt but not SPRTN-E112A cleaved CHK1 in at least three sites, generating three cleaved N-terminal fragments [Cleaved Products (CPs) 1, 2 and 3], as detected by Flag antibody (Fig. 5a). Similar N-terminal CHK1 fragments were obtained on endogenous or over-expressed CHK1 protein *in vivo* (Fig. 5b, c). Strikingly, these cleaved CHK1 products were completely absent in cells treated with three independent siRNA sequences targeting SPRTN (Fig. 5c). Altogether, these results further support the conclusion that CHK1 is a substrate of SPRTN *in vitro* and *in vivo*. As FA induces DPC formation and consequently increases SPRTN recruitment to chromatin (see also Fig. 6a-f) to proteolytically cleave chromatin-bound substrates, we wondered whether cleaved fragments of CHK1 are also enhanced in FA-treated cells (Fig. 5d and Supplementary Fig. 5 a). Indeed, increasing concentrations of FA stimulated the formation of cleaved N-terminal fragments of endogenous CHK1 (Fig. 5d) or over-expressed CHK1 (Supplementary Fig. 5 b), which mimic the SPRTN-dependent products of CHK1 cleavage *in vitro* and *in vivo.*

**Figure 5.**
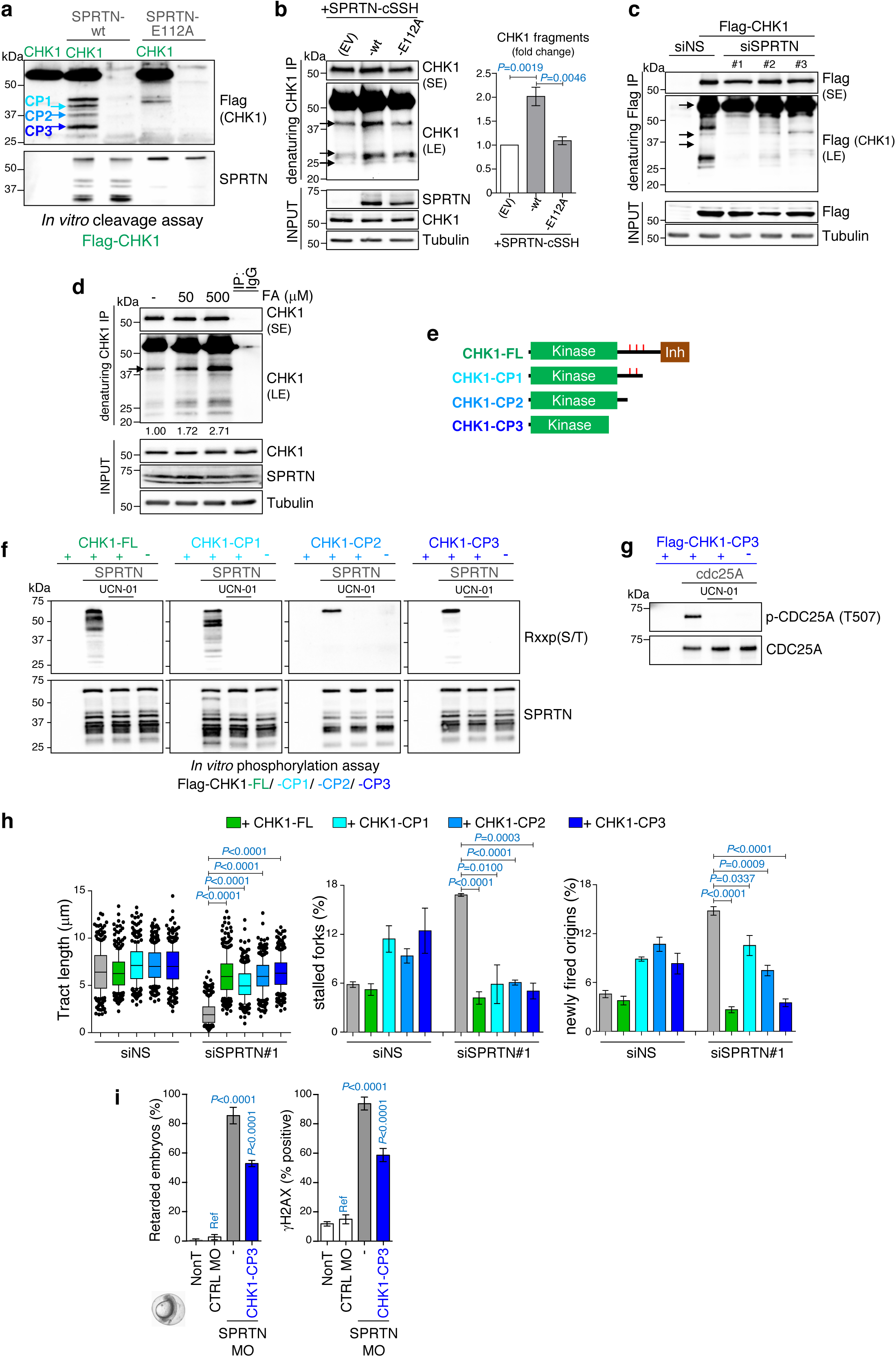
SPRTN cleaves CHK1 and releases kinase-active CHK1 fragments that are able to facilitate DNA replication and zebrafish development. **a**, SPRTN cleaves CHK1 *in vitro.* Purified Flag-CHK1 was incubated with SPRTN-wt or SPRTN-E112A (a protease-dead variant). Anti-Flag immunoblotting detected N-terminal CHK1 fragments released by the SPRTN proteolytic activity, termed CHK1 Cleavage Products (CPs) and indicated by blue arrows. Representative immunoblots from three replicates. **b**, SPRTN cleaves endogenous CHK1 *in vivo.* Whole cell extracts from HEK293 cells ectopically expressing SPRTN-wt or SPRTN-E112A were SDS-denatured and endogenous CHK1 was then immuno-purified using a CHK1 antibody raised against the N-terminal CHK1 epitope. **c**, SPRTN proficient cells exhibit CHK1 fragments that are absent in SPRTN-depleted cells. Whole cell extracts from HEK293 cells ectopically expressing Flag-CHK1 were SDS-denatured and CHK1 was then immuno-purified using anti Flag-beads. SE; shorter exposure, LE; lower exposure Representative immunoblots from three replicates. **d**, Addition of increasing concentrations of the DNA-protein crosslink inducer formaldehyde (FA, 1 h) correspondingly enhances the formation of endogenous CHK1 fragments in HEK293 cells. Numbers indicate the relative intensities of the arrowed band. Whole cell extracts were SDS-denatured and CHK1 purified with the CHK1 antibody raised against the N-terminal CHK1 epitop. LE: low exposure; HE: high exposure. Immunoblots shown represent three independent experiments. **e**, Graphical representation of the three N-terminal CHK1-CPs released by SPRTN activity. The approximate size of these CPs was estimated from the immunoblots and all contain the kinase domain. Based on these sizes, we made CHK1-CP1, CHK1-CP2 and CHK1-CP3 DNA constructs for the expression of these fragments in vertebrate systems encompassing CHK1 amino acids 1-338, 1-293 and 1-237, respectively. Red marks represent the location of Ser296, Ser317 and Ser345. Inh: autoinhibitory domain. **f**, *In vitro* phosphorylation assay showing that all three CHK1-CPs and CHK1-FL are kinase active and phosphorylates SPRTN. Flag-CHK1 variants were purified and incubated with SPRTN protein at 37°C for 2h. CHK1-mediated phosphorylation was detected with a specific antibody recognising the sequence R-x-x-p(S/T), a CHK1-phosphorylated substrate consensus sequence. **g**, CHK1-CP3 is also kinase active towards well-characterised CHK1 substrate CDC25A (0.5 μg, 2h) *in vitro.* Representative of three replicates. **h**, Ectopic expression of any CHK1-CPs, similar to expression of CHK1-FL (full-length), rescues the DNA replication defects in SPRTN-inactivated HEK293 cells (fork velocity, newly fired origins and stalled forks) assessed by DNA fiber assay analysis. Left: mean ± 25-75 percentile range (box) ± 1090 percentile range (whiskers); centre and right: mean ± SEM. > 100 DNA fibers were analysed per condition and experiment; n=3 experiments, two-tailed Students t-test. **i**, Ectopic expression of CHK1-CP3 in zebrafish embryos rescues the developmental defects and DNA damage elicited by SPRTN-deficiency. Mean ± SEM; n=3, two-tailed Student′s *t*-test.

We estimated the size of these fragments by their molecular weight and designed constructs mimicking these CPs for their expression in vertebrates (Fig. 5e). SPRTN-cleaved CHK1 fragments are kinase active as they were able to phosphorylate SPRTN or the well-characterised CHK1 substrate Cdc25A *in vitro* (Fig. 5f, g) ^49^.

Most notably, the N-terminal CHK1 products were fully functional *in vivo*, as their ectopic expression was able to restore DNA replication fork velocity and suppress new origin firing and fork stalling in SPRTN-depleted cells (Fig. 5h and Supplementary Fig. 5b). Importantly, the recovery of DNA replication velocity with the N-terminal CHK1 product CP3 was sensitive to the CHK1 kinase inhibitor UCN-01 but resistant to the ATR kinase inhibitor VE-821 (Supplementary Fig. 5c, d), and this rescue was completely dependent on the kinase active centre, located at aspartic acid 130 (D130) (Supplementary Fig. 5e). These results further suggest that the N-terminal CHK1 fragments, which mimic the cleaved CHK1 fragments generated by SPRTN proteolysis, are kinase active. Similarly, expression of the N-terminal CHK1 product CP3 corrected developmental defects and suppressed DNA damage in SPRTN-depleted zebrafish embryos to the same extent as full-length, wild-type CHK1 (Fig. 5i, compare with Fig. 2f, g).

### CHK1 stimulates recruitment of SPRTN to chromatin

Our results so far suggest that SPRTN is needed for proper activation of CHK1 during physiological DNA replication (Fig. 1). This CHK1 activation occurs by both (i) SPRTN proteolysis-dependent eviction of CHK1 from replicative chromatin (Fig. 4 g-k) and (ii) by simultaneous cleaving off the inhibitory/regulatory C-terminal parts of CHK1 (Fig. 5a-e). Paradoxically, ectopic expression of CHK1 restored various tested phenotypes of SPRTN-deficiency both in human cells and zebrafish embryos (Figs. 2, 5h, i and Supplementary Fig. 5b, d). As siRNA and MO-mediated SPRTN depletion is never 100% efficient, we speculated that ectopic CHK1 expression stimulates, by phosphorylation (Fig. 5f), the remaining SPRTN to process DPCs during DNA synthesis. Indeed, biochemical analysis demonstrated that SPRTN-inactivated cells still retain approximately 20-30% of SPRTN protein (Fig. 6a, b). To investigate this possible cross-talk between SPRTN and CHK1, we monitored SPRTN recruitment to chromatin after ectopic CHK1 expression (Fig. 6c, d). The amount of SPRTN on chromatin was substantially increased after CHK1 overexpression, suggesting that CHK1 induces SPRTN recruitment to chromatin. Importantly, the residual endogenous SPRTN in Δ-SPRTN cells (≈30% of control cells; Fig. 6a, b) hyper-accumulated on chromatin after CHK1 overexpression, almost to the same level as in untreated wild-type cells (Fig. 6c, d). To further support these results, we induced SPRTN-SSH expression in doxycycline-inducible HEK293 cells and ectopically coexpressed CHK1-wt or the N-terminal CHK1 cleaved fragments (CP1, 2 and 3). Ectopic expression of CHK1-wt stimulated SPRTN-SSH accumulation on chromatin. Strikingly, SPRTN-SSH recruitment to chromatin was even further enhanced after ectopic expression of the N-terminal CHK1 fragments (Fig. 6e, f). These results explain why the ectopic expression of CHK1-wt or its N-terminal fragments restores the phenotypes arising from SPRTN-deficiency in human cell lines and zebrafish embryos.

**Figure 6.**
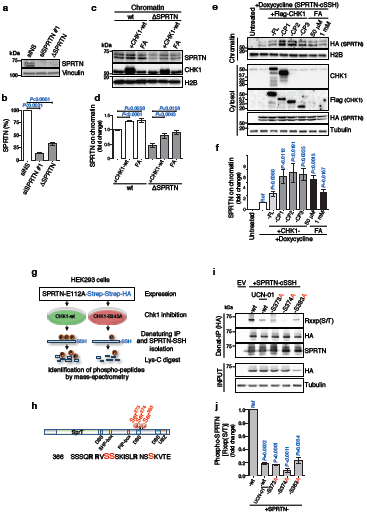
CHK1 phosphorylates SPRTN and regulates its recruitment to chromatin. **a-b**, siRNA SPRTN-depleted HEK293 cells or haploinsufficient SPRTN HeLa cells (ΔSPRTN) still contain a residual amount of SPRTN. Data are representative immunoblots and quantification of SPRTN normalised to vinculin. Mean ± SEM; n=3 experimental replicates, two-tailed Student′s *t*-test. **c-d**, CHK1 promotes SPRTN retention on chromatin. HeLa-wt or-ΔSPRTN cells ectopically expressing CHK1-wt, or treated with formaldehyde as a positive control, were subjected to cellular fractionation and endogenous SPRTN protein level was assessed in the chromatin fraction. Residual SPRTN in ΔSPRTN cells (approx. 30% of HeLa-wt cells) was also recruited to chromatin. Data are representative immunoblots and quantification of SPRTN on chromatin. Mean ± SEM; n=3 experimental replicates, two-tailed Student′s *t*-test. **e-f**, Overexpression of CHK1-full length (FL) or N-terminal products (CP1, 2 or 3) promote SPRTN retention on chromatin. Doxycycline inducible stable cells expressing SPRTN-wt-cSSH cells were used and fractionated into cytosol (control for total HA-SPRTN expression) and chromatin fractions. Data are representative immunoblots and quantification of HA (SPRTN) on chromatin. Mean ± SEM; n=3 experimental replicates, two-tailed Student′s *t*-test. **g**, Experimental strategy to identify the CHK1 phosphorylation sites on SPRTN by Mass-Spectrometry (see Methods). **h**, Diagram depicting human SPRTN with the CHK1 phosphorylation sites Ser373, Ser374 and Ser383 identified here. Known domains of SPRTN are also illustrated. SprT: protease domain; DBS: DNA binding site; SHP box: p97 interacting region; PIP box: PCNA interacting peptide; UBZ: ubiquitin binding domain. Sequence below (from Swissprot: Q9H040) shows the location of these sites highlighted in red and amino acids surrounding the region. **i-j**, The phosphorylation degree at a CHK1 consensus target sequence is diminished in the phospho-deficient mutants SPRTN-S373A, -S374A or -S383A. Different SPRTN variants were expressed in HEK293 cells and purified in denaturing conditions. An antibody recognizing the epitope Rxxp(S/T) (phospho-Ser/Thr preceded by Arg at position -3) was used to visualize CHK1 phosphorylated targets. The CHK1 inhibitor UCN-01 was used as a control. Data are representative immunoblots and quantification of CHK1 driven phosphorylation of SPRTN, normalised to total SPRTN protein level in denatured sample. Mean ± SEM; n=3 experimental replicates, two-tailed Student′s *t*-test.

### CHK1 phosphorylates the C-terminal regulatory part of SPRTN

To identify CHK1 phosphorylation sites (Fig. 5f) on SPRTN *in vivo*, we performed mass spectrometry. SPRTN-E112A-SSH was ectopically expressed in HEK293 cells co-transfected with either CHK1-wt or the kinase-defective CHK1-S345A, and anti-SSH precipitates were isolated under denaturing conditions (Fig. 6g, Supplementary Fig. 6a). The SPRTN protease dead mutant (E112A) was used to avoid SPRTN auto-cleavage activity ^20^ (Fig 5a, lower panel). Mass spectrometry analysis identified three main CHK1 phosphorylation sites on SPRTN - serines 373, 374 and 383 - (Fig. 6h and Supplementary Fig. 6b, c) located in the C terminal part of SPRTN. These CHK1 phosphorylation sites on SPRTN correspond to a consensus target motif of CHK1 (R/K-x-x-p(S/T) ^50^. To validate our results, we isolated SPRTN-wt-SSH or -S373A, -S374A and -S383A variants from HEK293 cells treated with DMSO or the CHK1 inhibitor UCN-01. SPRTN isolates were analysed by Western blot using an antibody that recognises CHK1 consensus sites (Fig. 6i, j). SPRTN-wt was phosphorylated in control cells but this phosphorylation signal was strongly decreased when cells were treated with UCN-01. Importantly, SPRTN phosphorylation was strongly reduced (S373A or S383A) or almost completely abolished (S374A) even when only one of the three identified serines (S373, S374 and S383) was mutated. This suggests that the C-terminal, regulatory part of SPRTN is phosphorylated by CHK1 at these specific serines under steady state conditions.

### SPRTN-CHK1 cross-activation loop

To biologically validate the biochemical evidence that CHK1 phosphorylates SPRTN and thus regulates SPRTN’s function, we depleted SPRTN in HEK293 cells and tested the effects of ectopically-expressing variants of SPRTN that either could or could not be phosphorylated by CHK1. SPRTN-wt and SPRTN-phospho-mimetic variants S373E and S383E restored DNA replication defects (Fig. 7a-c) and promoted CHK1 release from chromatin (Fig. 7d). Conversely, SPRTN phospho-defective variants S373A/374A and SPRTN-S383A completely failed to restore these phenotypes in SPRTN-depleted cells. Similar findings were observed in zebrafish embryos (Fig. 7e, f). These data suggest that CHK1-mediated phosphorylation of SPRTN is essential for SPRTN’s function *in vivo.*

**Figure 7.**
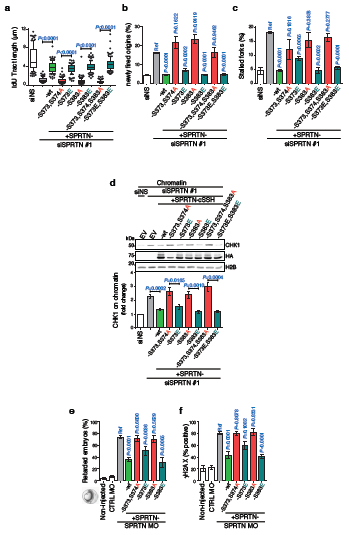
CHK1 targeted phosphorylation of SPRTN is essential for its biological function. **a-c** DNA replication analysis based on DNA fibers showing that SPRTN phospho-defective mutants are functionally impaired and unable to rescue DNA replication fork velocity (a), suppress new origin firing (b) and decrease stalled forks (c) in SPRTN-depleted HEK293 cells, whereas SPRTN phospho-mimetic mutants (S to E) are functional and recover a normal DNA replication phenotype. Left: mean ± 25-75 percentile range (box) ± 10-90 percentile range (whiskers); right: mean ± SEM. > 100 DNA fibers were analysed per condition and experiment; n=3 experiments, two-tailed Student′s *t*-test. **d**, SPRTN phospho-defective mutants failed to promote the eviction of CHK1 from chromatin fraction. Chromatin fractions from HEK293 cells were immunoblotted to show the effect of SPRTN variants on CHK1 release from chromatin. Data are representative immunoblots and quantification of CHK1 on chromatin fraction normalised to histone H2B. Mean ± SEM; n=3 experimental replicates, two-tailed Student′s *t*-test. **c** and **f**, SPRTN phospho-defective mutants were unable to rescue the zebrafish embryo developmental retardation (c) and DNA damage (d, γH2AX positive signal) induced by SPRTN deficiency. SPRTN-depleted embryo cells (SPRTN MO) were co-injected with plasmids for the expression of SPRTN variants. >70 embryos were scored per experiment. Mean ± SEM; n=3 independent experiments, two-tailed Student′s *t*-test.

To demonstrate our model in which a SPRTN protease-CHK1 kinase cross-activation loop processes DPCs in front of the DNA replication fork, we measured the progression of DNA replication fork over FA-induced DPCs by DNA fiber assay (Fig. 8a). Transient and low dose FA treatment strongly reduced DNA replication fork velocity, but this could be significantly reversed by expression of either CHKl-wt or SPRTN-wt. This demonstrates that both SPRTN and CHK1 prevent DPC-induced DNA replication fork slowing and stalling.

**Figure 8.**
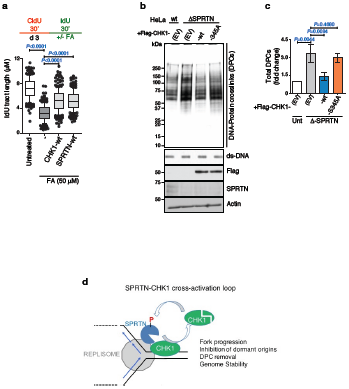
CHK1 and SPRTN promote replication fork progression by removing DPCs. **a**, Ectopic expression of CHK1-wt or of SPRTN-wt promotes replication fork progression over FA-induced DPCs in HEK293 cells. Upper panel shows the strategy for DNA fiber assay and lower graph plots the quantification of IdU tract length. >100 DNA fibers were analysed per condition and experiment. Mean ± 25-75 percentile range (box) ± 10-90 percentile range (whiskers) n=3 experiments, two-tailed Student′s *t*-test. **b-c**, Overexpression of CHK1-wt, but not of CHK1-S345A, reduces the amount of DPCs in HeLa-ΔSPRTN cells. DNA Protein Crosslink extracts were visualized by silver staining. ds-DNA blot is shown as a loading control. The immunoblots for Flag, SPRTN and actin were prepared from whole cell lysates. Data are representative immunoblots (b) and quantification (c) of CHK1 on chromatin fraction normalised to histone H2B. Mean ± SEM; n=3 experimental replicates, two-tailed Student′s /-test. **d**, Model for the SPRTN-CHK1 cross-activation loop. During steady-state DNA replication, SPRTN evicts CHK1 from replicative chromatin and thus SPRTN stimulates a functional steady-state CHK1 activation. Steady-state active CHK1 phosphorylates SPRTN on C-terminal serines. CHK1 activity enhances the recruitment of SPRTN to chromatin and its activity towards DPC proteolysis. This steady-state SPRTN-CHK1 cross-activation loop enable
s normal replication fork velocity, inhibits unscheduled origin firing, stabilises replication forks and removes DPCs under normal physiological conditions to safeguard genome stability.

As final confirmation that CHK1-overexpression indeed promotes the removal of DPCs in SPRTN-defective cells, we isolated total DPCs from wt and Δ-SPRTN Hela cells. A-SPRTN cells contain about 2-to 3-fold more DPCs than wt cells (Fig. 8b, c). However, ectopic expression of CHK1-wt, but not of kinase deficient variant CHK1-S345A, reduced the levels of DPCs in Δ-SPRTN cells to roughly the same level as in wt cells. Altogether, these results support our model (Fig. 8d) proposing that a SPRTN-CHK1 cross-activation loop works in DPC removal during physiological DNA replication to prevent DNA replication stress and preserve genomic stability.

## Discussion

Our results demonstrate the existence of an evolutionarily-conserved protease-kinase axis during physiological DNA replication that is essential for DPC proteolysis and prevention of DNA replication stress. We propose a model (Fig. 8d) wherein SPRTN protease and CHK1 kinase work in a cross-activation loop that operates during steady-state DNA synthesis. SPRTN protease activity releases CHK1 from replicating chromatin and this regulates DNA replication fork progression, suppression of dormant origin firing and prevents replication fork stalling, as well as the fine-tuning of CDK1/2 activity. These processes are essential for genome stability and embryonic development under physiological conditions as demonstrated in human cell lines and zebrafish embryos, respectively. In turn, CHK1 phosphorylates SPRTN at C-terminally located serines. SPRTN further accumulates on chromatin to proteolyse DPCs and thus allow unperturbed DNA synthesis. Our findings demonstrate that: (i) the SPRTN protease is tightly regulated and recruited to chromatin during DNA synthesis by CHK1 and at the same time (ii) SPRTN proteolysis evicts, cleaves and thus consequently activates CHK1 during steady-state DNA replication to fine-tune DNA replication and thus prevent DNA replication stress, one of the main causes of genome instability ^2,51^.

### Regulation of the SPRTN protease by CHK1 phosphorylation

The pleiotropic nature of SPRTN protease activity and its tight association with the replisome suggest that SPRTN must be tightly regulated. So far, the regulation of SPRTN has only been investigated upon exposure to genotoxic conditions such as UV light and FA. SPRTN accumulation at UV-induced DNA lesions strongly depends on its PCNA-binding PIP box and ubiquitin-binding UBZ domain ^52–54^. On the other hand, SPRTN activation in response to DPCs induced by 1 mM FA involves SPRTN deubiquitination, but does not depend on the PIP box and UBZ domains ^21^.

However, how SPRTN is regulated under physiological conditions is unclear. This is a critical question as SPRTN is an essential mammalian gene from early embryogenesis and is physically present at DNA replication forks and possesses pleotropic protease activity. It has been proposed that ssDNA promotes SPRTN-dependent substrate cleavage while dsDNA induces SPRTN auto-cleavage to negatively regulate SPRTN activity ^21^. There are two arguments against this model: (i) SPRTN is an essential enzyme for DPC repair during DNA replication, yet DPCs prevent the helicase-polymerase uncoupling and ssDNA formation that would be required to activate SPRTN-dependent substrate cleavage in this model, (ii) SPRTN protease activity towards DNA-binding substrates is also stimulated by dsDNA ^20,22^,

Our results suggest a model in which SPRTN protease activity is regulated by CHK1 kinase activity during physiological DNA replication. CHK1 constitutively phosphorylates SPRTN at the C-terminally located serines S373, S374 and S383. We show that CHK1-dependent SPRTN phosphorylation is essential for DNA replication and embryonic development under physiological conditions. We demonstrated that CHK1-dependent phosphorylation of SPRTN stimulates SPRTN recruitment to chromatin, and that these phosphorylation events are essential to regulate DNA replication, embryonic development and the release of CHK1 from chromatin. Further investigation should elucidate the phosphorylation dependant regulation of SPRTN.

### Regulation of CHK1 kinase during physiological DNA synthesis

It is well characterised how replication fork stalling and collapse leads to the formation of RPA-coated ssDNA and subsequent activation of CHK1 ^5^. However, a large body of evidence suggests that CHK1 is essential during unperturbed DNA synthesis when the ssDNA required for activating this canonical ssDNA-ATR-CHK1 signalling cascade is limited ^14,15,18,19^. Our data supports a novel mechanism for CHK1 chromatin release and activation during physiological DNA synthesis. We show that CHK1 associates with chromatin and components of DNA replication machinery and that SPRTN protease activity is required to release CHK1 from replicative chromatin during steady-state DNA synthesis.

We also demonstrated that SPRTN directly binds and cleaves the C-terminal part of CHK1 thus releasing at least three well detected the N-terminal CHK1 fragments (CP1, CP2 and CP3) *in vitro*. This effect is also observed with endogenous and ectopically expressed CHK1 *in vivo*. SPRTN protease cleavage of CHK1 generates active N-terminal CHK1 fragments that are able to phosphorylate SPRTN and restore DNA replication defects and developmental retardation in SPRTN-deficient human cells and zebrafish embryos, respectively.

The fact that CHK1 is a signalling molecule that activates downstream effectors but is mostly in an inactive state leads us to speculate that even a small amount of the CHK1 N-terminal fragments is sufficient to regulate physiological CHK1-signalling pathway activation during steady state DNA replication. In this way cells prevent the robust CHK1 activation usually observed after severe DNA damage that leads to apoptosis. Thus, we propose that SPRTN-dependent release and cleavage of even a small amount of CHK1 at/around DNA replication fork is sufficient to control DNA replication fork progression and ensure genome stability

Interestingly, the activation of CHK1 by cleavage of its C-terminal part was also observed in chicken DT40 and various human cells treated with Cisplatin and UV light as well as the well-known DPC-inducing agents, Campothecin and Etoposide ^55,56^. In some cases, this cleavage was caspase-dependent while in others it was not, pointing to the existence of a protease that cleaves and activates CHK1 ^55,56^. These reports and the herein described results that the N-terminal CHK1 kinase active fragments restore DNA replication and developmental defects in SPRTN-depleted cells and zebrafish embryos, respectevely (Supplementary Fig. 5 b, c) further support our model (Fig. 8d) that the SPRTN protease-CHK1 kinase cross-activation loop is operational during physiological DNA replication.

## Author contributions

S.H. and I.T performed the majority of the experiments. M.D.B and M.P. performed zebrafish experiments. M.Po. performed in vitro cleavage and interaction studies, J.F and D.P performed iPOND, B.V. performed DPC isolation, J.O. prepared Fig. 6a-b, D.L. analysed chromosomal aberrations and K.W. performed cell cycle analysis. A.N.S. created stable cell lines. I.T., I.V., R.F. performed mass-spectrometry and data analysis. S.H. initiated the project. S.H., I.T. and K.R. designed and analysed experiments. S.H., I.T., J.F. and K.R. prepared the final version of the manuscript. K.R. supervised the project.

## Acknowledgments

We thank Cornelia Donow for technical assistance and excellent fish care. This work was supported by Medical Research Council-UK (MC-EX-MR/K022830/1) and Oxford Cancer Research Centre to K.R. S.H. was supported by Goodger and Schorstein Scholarship University of Oxford and a shortterm postdoc fellowship by the European Institute of Innovation and Technology. The lab of MP is supported by Deutsche Forschungsgemeinschaft grant (PH144-4/-1) and the Boehringer Ingelheim Ulm University Biocenter. Mass spectrometry analysis was performed in the TDI MS Laboratory led by Benedikt M. Kessler. We thank E. Despras and P.L. Kannouche for providing us with ssDNA staining protocols, O. Gileadi and J. A. Newman for their help with SPRTN protein purification and G. Dianov for Flag-CHK1 construct. We thank T. Carr for critical reading of this manuscript.

## Supplementary Figure Legends

**Supplementary Figure 1.** Related to main figure 1.

**Analysis of cell cycle profile, CHK1 status and recruitment in chromatin, and CHK1-downstream signalling in SPRTN-deficient cells.**

**a-c**, Cell cycle analysis of HEK293 cells depleted of SPRTN with three different siRNAs. **a**, propidium iodide profiling. **b**, cell cycle profile from quantification of **a**. **c**, propidium iodide incorporation in cells in **a**. Data are representative of three replicates.

**d-e**, CHK1 was enriched on chromatin fraction (see also main figure 4a) in HEK293 cells depleted of SPRTN or treated with formaldehyde (FA; 50 μM, 1h) and its phosphorylation not enhanced. Cells were subjected to cellular fractionation to independently analyse chromatin fraction (upper panel of immunoblots) and phosphorylation in nuclear soluble and cytosolic fractions (medium and lower panels). CHK1 on chromatin was quantified. Mean ± SD; n=3 replicates, two-tailed Student′s /-test.

**f**, CDC25A total protein level is stabilized in SPRTN-depleted HEK293 cells. Data are quantifications of immunoblots from Main Fig. 1f. Mean ± SD; n=3 replicates, two-tailed Student′s /-test.

**g-h**, phospho-CDK substrates are enhanced in SPRTN-depleted HEK293 cells. Data are representative immunoblots of whole cell lysates using a specific antibody that recognizes phosphorylated CDK substrates, and quantification normalised to actin. Mean ± SD; n=3 replicates, two-tailed Student′s /-test.

**Supplementary Figure 2.** Related to main figure 2.

**a**, Immunoblots showing the expression and phosphorylation degree of different CHK1 variants overexpressed in HEK293 cells (upper panel), and cell cycle profile of HEK293 cells as indicated (lower panel). Related to main figure 2a.

**b**, Immunoblots showing the efficiency of SPRTN depletion alone or combined with the expression of CHK1-wt. Related to main figure 2c.

**c**, Graphical illustration of embryos developing normally after 6, 8 and 10 hpf (upper row), and graphical illustrations of embryos at 10 hpf at the indicated conditions (lower row) as observed in Fig. 2d.

**Supplementary Figure 3.** Related to main figure 3.

**SPRTN-deficient cells retain the ability to form ssDNA after HU- or CPT-induced replication stress or DNA double strand formation, respectively. a-b**, Data shown are representative immunofluorescent microscopy images of BrdU foci (marker for ssDNA) in S-phase (marked by CldU positive cells) HeLa cells treated with either siNS or siSPRTN and quantification. >70 S-phase cells were scored per condition per experiments. Mean ± SEM, n=3 experimental replicates, twotailed Student′s *t*-test.

**Supplementary Figure 4.** Related to main figure 4.

**SPRTN protease evicts CHK1 kinase from chromatin**

**a**, Immunoblot analysis of chromatin fractions from control or SPRTN-depleted HEK293 cells cotransfected with empty vector (EV), SPRTN-WT or SPRTN-Y117C (patient variant with decreased protease activity), showing eviction of CHK1 from chromatin. pRPA2 and DNA damage (gH2AX) are shown as markers for DNA damage. HU (1 mM, 1 h) was used as a positive control.

**b**, Quantification of CHK1 on chromatin, normalised to histone H2B. Mean ± SD, n = 5 experiments, two-tailed Student′s *t*-test.

**Supplementary Figure 5.** Related to main figure 5.

**The N-terminal fragments of CHK1 (CP1 or CP3) restore DNA replication defects in SPRTN-deficient cells.**

**a**, Addition of increasing concentrations of the DNA-protein crosslink inducer formaldehyde (FA, 1 h) correspondingly enhances the formation of Flag-CHK1 fragments in HEK293 cells co-transfected with Flag-CHK1. Numbers indicate the relative intensities of the arrowed band. Whole cell extracts were SDS-denatured and Flag-CHK1 purified with the Flag antibody.

**b**, Expression of Flag-CHK1 variants in control and SPRTN-depleted cells (the Flag tag is located at the N-terminus). Anti-CHK1 antibody could not detect CHK1-CP2 and CHK1-CP3 due to loss of epitopes, but anti-Flag antibody could. Related to main figures 5d-g.

**c**, Analysis of DNA replication for velocity by DNA fiber assay. The ATR inhibitor VE-821 (5 μM) or the CHK1 inhibitor UCN-01 (300 nM) severely slow DNA replication fork velocity in control cells (siNS). Ectopic expression of the smallest N-terminal CHK1 fragment (CP3) rescues DNA replication fork velocity in SPRTN-depleted cells (siSPRTN). The rescue effect of CHK1-CP3 fragment on DNA replication fork velocity in SPRTN-depleted cells is CHK1 kinase dependent (sensitive to UCN-01) but ATR kinase independent (resistant to VE-821). Upper diagram shows the experimental strategy including DNA fiber assay (d: day). Graph below shows the data for replication fork velocity (mean ± 25-75 percentile range (box) and 10-90 percentile range (whiskers)). > 100 DNA fibers were analysed per condition and experiment; n=3 experiments, two-tailed Student′s *t*-test. Related to main figure 5g.

**d**, Immunoblot analysis for b shows the efficiency of SPRTN depletion and the expression of Flag-CHK1-CP3 in the presence of the CHK1 inhibitor UCN-01 or the ATR inhibitor VE-821.

**e**, The active kinase domain of the N-terminal CHK1 fragments (CP1 and CP3) is required for the rescue of DNA replication defects in SPRTN-depleted HEK293 cells. D130A: kinase dead CHK1 variants with a mutation in Aspartic acid (D) 130 to Alanine (A) located in the catalytic domain. Data shows DNA fiber assay analysis showing fork tract length (mean ± 25-75 percentile range (box) and 10-90 percentile range (whiskers)). > 100 DNA fibers were analysed per condition and experiment; n=3 experiments, two-tailed Student′s *t*-test. Related to main figure 5g.

**Supplementary Figure 6.** Related to main Fig. 6.

**Identification of CHK1 phospho-targets in SPRTN by Mass-Spectrometry.**

**a**, Samples for Mass-Spectrometry analysis were prepared by co-expression of the protease-dead mutant SPRTN-E112A (to prevent autocleavage) tagged with Strep-Strep-HA and either with CHK1-wt or the CHK1-S345A mutant, followed by denaturing-IP isolation of SPRTN.

**b**, Example of MS/MS fragmentation spectra (PEAKS; intensity up to 24%) of SPRTN peptide 362-376 (K.NSVSSSSQRRVSSSK.I) detected in the CHK1-WT overexpression sample, showing a phosphorylation in amino acid S373. Arrows point to the b and y ions which strongly support that the phosphorylation site is on S373 (p-site localization score of 8.18).

**c**, Extracted ion chromatograms (XIC; 4ppm) for SPRTN 362-376 peptide (top panel) and for SPRTN 377–384 peptide (bottom panel) containing amino acids S373, 374 and S383, respectively, comparing peptides species detected when CHK1-wt (left) or CHK1-S345 (right) was overexpressed. Each red tone corresponds to different peptides species containing potential phosphorylation at S373, 374 or S383, whereas green tones correspond to non-phosphorylated peptides. PTMs were automatically assigned by PEAKS software. Pie charts areas represent peptide intensities. *: Peptide identification performed by matching m/z and retention time from existent data. Not ID = not identified. + coeluting peptides with almost identical m/z and retention time but with different p-site assignment.

**d**, DNA replication analysis based on DNA fibers showing that SPRTN single phospho-defective mutants (S to A) are functionally impaired and unable to rescue DNA replication, whereas SPRTN phospho-defective mutants (S to E) are functional. Related to main Fig. 6e.

## Methods

Key resources used in this study are listed and described in Supplementary Table 2. Oligonucleotide sequences are listed in Supplementary Table 3.

### Experimental models and cell culture

U2OS, HeLa and HEK293 human cells were maintained in DMEM (Sigma-Aldrich) supplemented with 10% FBS (Gibco) and 100 I.U./mL penicillin / 0.1 mg/mL streptomycin at 37°C in a humidified incubator with 5% CO2, and tested for mycoplasma contamination. CRISPR partial knockout Δ-SPRTN HeLa cells^20^ were maintained as above. Media for doxycycline-inducible stable HEK293 Flp-In TRex SPRTN-wt or SPRTN-Y117A cell lines^20^ was additionally supplemented with 15 μg/ml blasticidin / 100 μg/ml hygromycin. Wild-type zebrafish (*Danio rerio* EK and AB strains) were kept under standardized conditions in a circulating water system on a 14 hours light and 10 hours dark cycle. All procedures involving zebrafish embryos adhered to current European regulations for the use of animals and were approved by the local authorities of Ulm University.

### Bacterial strains

Competent *E. coli* DH5alpha (Invitrogen) and *E. coli* Rosetta2 (Merck; Novagen), used for cloning/ plasmid amplification or for protein expression, respectively, were transformed with plasmid and grown in LB medium at 37 °C.

### Subcloning and site directed mutagenesis

The pCINeo/CMV-Flag-CHK1-wt plasmid^11^ was used for expression in mammalian cells and to generate all CHK1 variants by site-directed mutagenesis. For the expression of CHK1 in Zebrafish, CHK1 variants were subcloned into pCS2-GFP vector by PCR from previous plasmid and EcoRI/XbaI insertion. Site-directed mutagenesis was performed by plasmid PCR using specific mutagenic primers (Supplementary Table 3) designed by the quick-change primer design software (Invitrogen) and AccuPrime Pfx DNA-polymerase (Invitrogen). Mutations were then verified by sequencing at Source BioScience, Oxford, UK.

### RNA interference and Plasmid transfection

siRNA transfections were performed using Lipofectamine RNAiMax reagent (Invitrogen) according to manufacturer’sinstructions. Depletion was assayed 72 h post-transfection. Plasmid transfections were performed using either FuGene HD reagent (Promega) following manufacturer’s instructions or by PEI method. Protein expression was assayed 12-48 h post-transfection.

### Denatured whole cell extracts

Cells were collected, washed three times in ice-cold PBS, and lysed in RIPA buffer (50 mM Tris-HCl, pH 7.4, 150 mM NaCl, 1 % NP-40, 0.5 % SDS, 0.1 % SDS, 3 mM EDTA, 20 mM N-ethylmaleimide (NEM), 1 mM DTT, protease inhibitors [1 μg/mL Chymostatin, 1 μg/mL Aprotinin, 1 μg/mL Leupeptin, 1 mM PMSF] and phosphatase inhibitors [1 mM Na_3_VO_4_x2H_2_O, 1 mM Na_4_P_2_O_7_, 20 mM NaF]). Extracts were then sonicated (20 cycles; 30/30 sec on/off) in a Bioruptor Plus sonicator (Diagenode).

### Cell fractionation

Cells were incubated in 2x volumes of Buffer A (10 mM HEPES, pH 7.4; 10 mM KCl; 1.5 mM MgCl2; 340 mM Sucrose; 10% Glycerol; 1 mM DTT; 5 mM EDTA; 20 mM NEM; protease and phosphatase inhibitors and 0.1% Triton-X100) on ice for 10 minutes. Sample was spun (350g, 4 min, 4 °C) and the supernatant was collected as the cytosolic fraction. Nuclei were then washed once with Buffer A without Triton-X100 and burst in 2x volumes of hypotonic Buffer B (3 mM EDTA; 0.2 mM EGTA; 1 mM DTT; 10 mM NEM; protease and phosphatase inhibitors) on ice for 5 min. NaCl to 250 mM and NP-40 to 0.6% were added and sample rotated for 10 min. After centrifugation at 2,500g for 5 min, the supernatant was collected as the nuclear soluble fraction. The pellet was washed twice with Buffer B with 0.6% NP-40 and 250 mM (first wash) or 150 mM (second wash) NaCl, and this was considered as the chromatin fraction. For chromatin nuclease extracts, chromatin fraction was washed in Benzonase buffer (50 mM Tris pH 7.5; 20 mM NaCl; 5 mM KCl; 3 mM MgCl_2_; protease and phosphatase inhibitors) and incubated in same buffer at 4°C with 100 u/mL Benzonase (Millipore).

### Denaturing immunoprecipitation

Denatured extracts were first obtained by lysing cells expressing tagged protein in denaturing buffer (50 mM Tris, pH 7.4; 137 mM NaCl; 5 mM KCl; 1% SDS; 1 mM DTT; 5 mM EDTA; 20 mM NEM; protease and phosphatase inhibitors), heating at 95°C for 5 min, shearing chromatin by 10-15 passes through a 22-gauge needle and sonicating –20 cycles; 30/30 sec on/off– in a Bioruptor Plus sonicator (Diagenode), before been clarified by centrifugation at 15,000g for 5 min. 10% of supernatant was retrieved for input. Denatured extract was diluted 10 fold in IP buffer (50 mM Tris, pH 7.4; 150 mM NaCl; 10 mM NEM; protease and phosphatase inhibitors) with 1% Triton X-100 to quench SDS, and precleared with blank sepharose (IBA) and 2 U avidin for 1 h. After incubation for 4 h to overnight with Strep-Tactin sepharose (IBA) or Anti-Flag M2 affinity beads (Sigma-Aldrich), extract was washed twice with IP buffer containing 500 mM NaCl and 1% Triton, and 3 times with IP buffer containing 137 mM NaCl and no detergents. Samples were finally eluted in 2x Laemmli buffer for 10 min at 95°C.

### Protein Co-immunoprecipitation

To isolate SPRTN-interacting proteins, non-denaturing whole cell lysates were prepared from corresponding HEK293 Flp-In TRex cell lines upon doxycycline induction of SPRTN-WT-cSSH (Strep-Strep-HA tag). In addition, cell lysates were also prepared from HEK293 cells transiently transfected with SPRTN-WT-cSSH and SPRTN-Y117-cSSH in order to compare the capturing efficiency between SPRTN-WT and SPRTN-Y117C (patient mutation), respectively. Cells were collected and lysed in non-denaturing lysis buffer [50 mM Tris pH 7.4; 150 mM NaCl; 1% Triton; 1mM EDTA; 10 mM NEM; protease and phosphatase inhibitors] with 2 U avidin (IBA Cat# 2-0204-015) at 4°C. Following centrifugation at 16,000 g for 10 min, the supernatant was kept at 4°C and the pellet was digested with Benzonase nuclease in Benzonase buffer (see above) at 37° C for one hour. Both supernatant and Benzonase digested fractions were pooled and 10% of this pooled fraction was stored at −20° C for later use as input. The rest of the lysate was diluted 10 times in IP buffer [50 mM Tris, pH 8; 150 mM NaCl; 10 mM NEM; protease and phosphatase inhibitors] and precleared with blank sepharose (IBA GmbH) for 1 hour at 4°C on a rotating wheel. Precleared samples were then incubated with Strep-Tactin sepharose beads (IBA Cat# 2-1201-010) for 2 hours at 4°C, washed five times in IP buffer containing 0.05% NP-40 and eluted in 2x Laemmli buffer (62.5 mM Tris-HCl, pH 6. 8; 2% SDS, 10% Glycerol, 2.5% β-mercaptoethanol; 0.001% Bromophenol blue) for 10 minutes at 95°C.

### Western blotting

Samples for WB were prepared following standard methods. Briefly, protein concentration in cleared protein extracts was measured by Lowry (Bio-Rad) and samples were denatured in Laemmli buffer (95°C, 5 min). SDS-PAGE was performed in TGS buffer. After transfer, membranes were blocked in TBST buffer with 5% milk (RT, 1 h), washed with TBST and incubated (overnight, 4 °C) with primary antibody (Supplementary Table 2) diluted in 2% BSA in TBST. Membranes were then washed four times in TBST, incubated (RT, 1 h) with HRP-conjugated secondary antibody in TBST with 1% milk, and washed 4 times in TBST. Protein was then visualized using ECL substrates (Thermo Scientific and Bio-Rad) in a ChemiDoc MP system (Bio-Rad) or with X-ray film. Quantification of immunoblots was performed using ImageLab software (BioRad) or ImageJ. Specific protein level values were calculated relative to the corresponding loading control. Statistical significances of replicate measurements were analyzed in GraphPad Prism.

### Isolation of Proteins On Nascent DNA: IPOND

iPOND was performed as described earlier^20^. Briefly, newly synthesized DNA (~2 × 10^8^ cells per condition) was labelled via incubation with 10 μM EdU for 10 min (in SPRTN-wt cells) or 15 min (in SPRTN depleted cells). For mature chromatin, cells were chased with thymidine for further 5 min. Chromatin fragmentation into 300-500 bp fragments was done in a Bioruptor Plus sonicator (Diagenode) (70cycles, 30/30 sec on/off). After conjugation of biotin to EdU by click chemistry (Dungrawala et at, 2015), proteins on EdU-labelled DNA were isolated with streptavidin-coupled agarose beads.

### DNA fiber assay

The DNA fiber assay was performed as described previously^20,25^. Asynchronous cells were labelled with 30 μM CldU for 30 min and then with 250 μM IdU for additional 30 min. For assay of genotoxic stress, IdU was incubated in the presence drugs. DNA replication was inhibited by with ice-cold PBS. Cells were lysed in 200 mM Tris-HCl pH 7.4, 50 mM EDTA and 0.5% SDS; DNA fibres were spread onto positively charged glass slides, fixed with 3:1 methanol: acetic acid (v/v), denatured with 2.5 N HCl, blocked with 2% BSA in PBS containing 0.001% Tween-20, and stained for 1 h at 37°C with anti-BrdU that specifically recognize either CldU (clone BU1/75(ICR1) or IdU (clone B44). After incubation, slides were washed three times with PBS containing 0.05% Tween-20 (PBS-T) and briefly blocked again. Anti-rat Cy3 (Jackson Immuno Research) and anti-mouse Alexa-Fluor 488 (Molecular Probes) were the respective secondary antibodies. Slides were washed 3 times with PBS-T, air-dried at RT and mounted with ProLong Gold antifade reagent (ThermoFisher). Microscopy was performed using a Leica DMRB microscope with a DFC360FX camera. Quantification of CldU- and IdU-labelled DNA tract lengths was done with ImageJ software on at least 100 fibers per condition in at least three independent experiments, converting arbitrary units to microns based on the microscope scale. Resulting “Tract length” values were represented in box graphs showing the mean (bar) with the 25-75 percentile range (box) and the 10-90 percentile range (whiskers) and statistically analyzed using GraphPad Prism. Analysis of newly fired origins and of stalled forks were performed as above but on at least 400 fibers per condition tested, and data was represented as plot the mean ± SEM of at least three independent experiments.

### Metaphase spreads

Metaphase spreads were performed as described previously^25^. HEK293 were treated with 100 ng/mL colcemid for 2 h, trypsinized, collected, and incubated with 0.4% KCl at 37 °C for 15 min. Following addition of fixative solution (3:1 methanol:acetic acid), cells were centrifugated at 1,000g for 10 min, resuspended in 5 mL fixative solution, incubated at RT for 15 min, centrifuged, resuspended in 1 mL 100% acetic acid, incubated at RT for 10 min, centrifuged, and finally resuspended in 5 mL of fixative solution. Metaphase spreads were prepared by dropping the cell suspension from a height onto pre-wetted slides. The slides were air dried, stained with DAPI and examined by microscopy. Analysis were performed on at least 30 metaphases per sample.

### Immunofluorescence Microscopy

Cells grown on round coverslips were washed once with ice cold PBS, pre-extracted for 3 mins with ice-cold pre-extraction buffer (25 mM HEPES pH 7.5; 50 mM NaCl; 1 mM EDTA; 3 mM MgCl_2_; 300 mM Sucrose; 0.5% Triton-X100) and fixed by adding methanol and 30 min incubation at −20 °C. Fixed cells were then washed three times with PBS, blocked with 3% BSA in PBS with 0.5% Tween-20 for 1 h at RT, incubated (90 min, RT) with the primary antibody, washed three times, and incubated with the corresponding secondary antibody (45 min, RT). Incubation with DAPI was done for 15 min at RT. Finally, coverslips were mounted onto clean glass slides with mounting media. Microscopy was performed using a Leica DMRB microscope with a DFC360FX camera.

### ssDNA foci

Detection of single-stranded DNA (ssDNA) under non-denaturing condition was done as described previously^57^ with minor modifications. 24 h after siRNA transfection, cells growing on coverslips were treated with 10 μM BrdU (Sigma-Aldrich) for further 48 h, and then pulse-labelled with 100 μM CldU for 30 min before been treated with replication interfering drugs. Cells were washed with PBS, pre-extracted, fixed and blocked as above. ssDNA was marked with anti-BrdU (clone B44; 1:100 dilution) for 1 h, then three washes with PBS, followed by secondary anti-mouse Alexa-Fluor 488 antibody (Molecular Probes, 1:1,000 dilution) for 30 min and three washes. The primary and secondary antibody conjugates were then fixed with 4% formaldehyde in PBS for 15 min followed by another wash. DNA was then denatured in 1.5 N HCl for 30 min and washed with PBS. For the detection of replication foci after denaturation, sample was incubated with anti-CldU antibody (clone BU1, 1:100 dilution) for 1 h, washed twice with PBS, then with high salt buffer (PBS with 200 mM NaCl, 0.2% Tween-20 and 0.2% NP-40) for 15 min and then with PBS again, then incubated with anti-rat IgG Cy3 antibody (Jackson Immuno Research, 1:2,000 dilution), and washed again. Finally, coverslips were mounted in mounting medium supplemented with 2x DAPI. Images were taken with a Leica DMRB microscope with a DFC360FX camera.

### DNA Protein Crosslink isolation and detection

DNA Protein Crosslinks (DPCs) were isolated using a modified Rapid Approach to DNA Adduct Recovery (RADAR) assay^58^ and as described previously^20^. Briefly, 1-2×10^6^ cells were lysed in 1 mL of buffer containing 6 M guanidine thiocyanate (GTC); 10 mM Tris–HCl, pH 6.8; 20 mM EDTA; 4% Triton-X100; 1% Sarkosyl and 1% DTT. DNA was precipitated by adding 100% ethanol, washed three times in wash buffer (20 mM Tris-HCl, pH 6.8; 150 mM NaCl and 50% ethanol). DNA was then solubilized in 1 mL of 8 mM NaOH. The DNA concentration was quantified by treating a small aliquot of DNA with proteinase K (Invitrogen) for 3 h at 50°C, followed by detection with PicoGreen dye (Invitrogen) according to manufacturer’s instructions. Equal dsDNA loading was confirmed by slot-blot immunodetection with an anti-dsDNA antibody (Abcam). Total DPCs after electrophoretic separation on polyacrylamide gels were visualized by silver staining using the ProteoSilver Plus Silver Stain Kit (Sigma-Aldrich) and specific proteins were detected by Western-blot.

#### SPRTN purification

SPRTN protein purification was performed as described previously^20^. *E. coli* was resuspended in lysis buffer (100 mM HEPES, pH 7.5; 500 mM NaCl; 10% glycerol; 10 mM imidazole; 1 mM Tris(2-carboxyethyl)phosphine (TCEP); 0.1% dodecylmaltoside (DDM); 1 mM MgCl_2_ and protease inhibitor cocktail (Merck)) and sonicated. 1 U Benzonase nuclease was added to lysates before cell debris was pelleted by centrifugation. Lysates were applied to a Ni-sepharose IMAC gravity flow column, washed with two column volumes of wash buffer (50 mM HEPES, pH 7.5; 500 mM NaCl; 10% glycerol; 45 mM imidazole; 1 mM TCEP), and eluted in elution buffer (wash buffer but with 300 mM imidazole). Elution fractions were applied directly to a 5mL Hitrap SP HP column (GE healthcare), washed with wash buffer 2 (50 mM HEPES, pH 7.5; 500 mM NaCl; 1 mM TCEP) and eluted with elution buffer (wash buffer but with 1 M NaCl). The purification tag was cleaved by the addition of 1:20 mass ratio of His-tagged TEV protease during overnight dialysis into buffer A (20 mM HEPES, pH 7.5, 500 mM NaCl, 0.5 mM TCEP). Samples were concentrated by ultrafiltration using a 30 kDa molecular weight cut-off centrifugal concentrator and loaded onto size exclusion chromatography using a HiLoad 16/60 Superdex 200 column (GE Healthcare) at 1 mL/min in buffer A. Protein identities were verified by LC/ESI-TOF Mass spectrometry^20^ and protein concentrations were determined by absorbance at 280 nm (Nanodrop) using the calculated molecular mass and extinction coefficients.

#### In vitro cleavage of CHK1

Recombinant SPRTN was purified from *E. coli.* Flag-CHK1 was purified from ectopically expressing HEK cells in non-denaturing conditions by immobilization in Flag-M2 affinity agarose beads and several washes containing either 0.5% Triton X-100 or 1 M NaCl, and then eluted with a Flag peptide (Sigma-Aldrich). Cleavage reaction was performed typically in 15 μL volume containing Flag-CHK1, 2 ug SPRTN, and a 100 bp dsDNA oligonucleotide probe (Supplementary Table 3; 30:1 molar ratio SPRTN:dsDNA) in cleavage buffer (25 mM Tris, pH 7.4; 150 mM NaCl) at 37°C for 4-8 h, and stopped by the addition of Laemmli buffer.

#### *In vitro* CHK1 kinase assay

Flag-CHK1 species (full length or cleavage products) were obtained from 10^8^ HEK cells previously transfected with expressing vectors and purified as above. For phosphorylation activity, purified Flag-CHK1 was incubated (37 °C, 2 h) with a substrate (SPRTN: 2 μg; cdc25A: 500 ng) in a kinase reaction buffer (10 mM Hepes, pH 7.4; 0.5 mM ATP; 10 mM MgCl_2_; 5 mM KCl; 0.1 mM DTT; 20 mM β-glycerophosphate and phosphatase inhibitors) in a total volume of 25 μL. The reaction was stopped by the addition of 5x Laemmli buffer and boiling.

### Manipulation and analysis of Zebrafish embryos

Wild-type zebrafish (EK and AB strains) were kept under standardised conditions in a circulating water system (Tecniplast, Germany). Eggs were collected from natural matings. At the one to two cell stage, fertilized eggs were microinjected with a previously described SPRTN MO^25^ and with *in vitro* synthesized, capped RNAs encoding different CHK1 variants fused to GFP, which were prepared from linearized plasmids using the mMessage mMachine SP6 Kit (Ambion). Alternatively, capped RNA encoding SPRTN variants were co-injected. Growth retardation was assessed at 9 hours post fertilization (hpf; 90% epiboly stage under control conditions) using an upright brightfield M125 microscope equipped with a IC80 HD camera (both Leica). Immunofluorescence stainings were essentially prepared as previously described^59^ using a rabbit-anti-gH2Ax antibody (Genetex, 1:200 dilution) and imaged at a M205FA microscope equipped with a DFC365FX camera (both Leica).

### Mass Spectrometry

Purified SPRTN was isolated by denaturing IP using Strep-Tactin beads (IBA), from 2.5×10^8^ HEK293 cells ectopically coexpressing SPRTN-E112A-SSH and either CHK1-WT or CHK1-S345A. SPRTN isolates were reduced with 5 mM DTT (Sigma-Aldrich; room temperature, 1 h), alkylated with 20 mM iodoacetamide (Sigma-Aldrich; room temperature, 1 h), and extracted using a double methanol/chloroform precipitation. Protein precipitates were resuspended in 6 M urea (with sonication), and diluted in a buffer containing 25 mM Tris-HCl (pH 8.5) and 1 mM EDTA until the urea concentration was < 1 M. Protein was subsequently digested with rLysC (Promega) for overnight at 37°C. Peptides were then desalted using a SOLA HRP SPE cartridge (Thermo Scientific Cat# 60109-001) and dried.

### Mass Spectrometry

Dried tryptic peptides were reconstituted in 12 μL of LC-MS grade water containing 2% acetonitrile and 0.1% trifluoroacetic acid. Samples were subsequently analyzed by nano-LC-MS/MS using a Dionex Ultimate 3000 UPLC coupled to a QExactive instrument (Thermo Scientific, Bremen, Germany) as described previously^60–62^. Peptides were online desalted on a PepMapC18 column (300 μm × 5 mm, 5 μm particle size, Thermo Fischer) for 1 min at a flow rate of 20 μL/min and separated on a directly coupled nEASY column (PepMap C18, 75 μm × 500 mm, 2 μm particle, Thermo Fischer) using a multistep gradient starting with 3 min at 2% Acetonitrile in 5% DMSO with 0.1% Formic acid (buffer B), followed by 60 mins linear gradient up to 35% buffer B at 250 nL/min flow rate, 7 min linear gradient up to 99% buffer B and maintained at 99% buffer B for 5 min at a flow rate of 250 nL/min before reverting to 2% buffer B for 3 min prior to a last 4 min at 99% solvent B. Full MS scans were acquired over the m/z range of 380—1800 at a resolution of 70,000 at 200 m/z (AGC target of 3×10^6^ ions). MS/MS data was acquired in a data dependent manner by selecting the 15 most abundant precursor ions for HCD fragmentation (CE of 28) and on MS/MS resolution of 17,500.

### Mass Spectrometry data analysis

LC-MS/MS data was analyzed by PEAKS (V7.5; Bioinformatics Solutions Inc, Waterloo, ON, Canada) using PEAKS database and PTM function. Data was searched against Human UniProtSwissprot database (downloaded on August 2016) enabling the decoy function whilst selecting LysC as enzyme (two missed cleavages; none non-specific cleavage), HCD as fragmentation source, mass tolerance of 10 ppm for precursor ions (MS data) and 0.05 Da for peptide fragment ions (MS/MS data); carboamidomethylation (C) as a fixed modification, and Oxidation (M), deamidation (NQ) and phosphorylation (STY) as variable modifications. PEAKS-PTM results were filtered using a 1% FDR at peptide level. Extracted ion chromatogram (XIC) of peptides of interest was performed using Xcalibur Qual Browser (Thermo Scientific) enabling Gaussian peak smoothing at 3 points and a mass tolerance of precursor ion (MS1) of 4 ppm. Peaks areas were automatically calculated by the software. Mass spectrometry data analysis for phospho-sites detection performed by PEAKS software is contained in Supplementary Table 1.

### Statistics and reproducibility

Every experiment was reproduced at least three times, with similar results, except for the experiments shown in Figs 2g and 7e-f, which were repeated twice. Unless otherwise stated, the statistical method used for comparison between experimental groups was unpaired two-tailed Student′s *t*-test carried out using GraphPad Prism. Statistical significance was expressed as a *P* value, where P<0.05 was considered a statistically significant difference.

## Data availability

The Mass Spectrometry raw data reported in this paper have been deposited in the ProteomeXchange Consortium via the PRIDE^63^ partner repository with the dataset identifier PXD006741.

### Reviewer account details

**Password:** ktPFQ6Oy

### Source data

DNA fibers, immunoblot quantifications,IF foci, etc

All other data supporting the findings are available from the corresponding author at a reasonable request.

